# KIR3DL1 and Tox identify clonally expanded encephalitogenic neuron-specific CD8+ T cells in autoimmune encephalitis

**DOI:** 10.1101/2024.03.25.586688

**Authors:** Sylvain Perriot, Samuel Jones, Raphaël Genolet, Amandine Mathias, Helen Lindsay, Sara Bobisse, Giovanni Di Liberto, Mathieu Canales, Lise Queiroz, Christophe Sauvage, Ingrid Wagner, Larise Oberholster, Marie Gimenez, Diane Bégarie, Marie Théaudin, Caroline Pot, Doron Merkler, Raphaël Gottardo, Alexandre Harari, Renaud Du Pasquier

## Abstract

Autoreactive CD8+ T cells are the principal suspects in autoimmune encephalitis (AIE) with antibodies targeting intracellular neuronal antigens So far, the search for neuron-autoreactive CD8+ T cells has been focused on a few autoantigens and did not yield convincing results. Here, we leveraged natural antigen presentation by hiPSC-derived neurons to look at the global autoreactive CD8+ T cell response, independently of pre-conceived hypothesis of the autoantigens involved in the disease. This unbiased approach allowed for the identification of rare polyclonal neuron-reactive CD8+ T cells in healthy donors, and contrastingly, expanded clonotypes in two patients with anti-Ri AIE. Detailed *ex vivo* phenotypic characterization of these clonotypes revealed a specific transcriptional program suggestive of a pathogenic potential. In particular, this subset can be identified by the expression of KIR3DL1 and TOX. Strikingly, we could also demonstrate that CD8+ T cells found in the brain of an anti-Ri AIE patient display a similar phenotype associated with cytotoxicity and encephalitogenic features.

## Introduction

Autoimmune encephalitis (AIE) are a heterogeneous group of neurological syndromes characterized by the presence of autoreactive antibodies targeting neuronal or glial antigens located in the intracellular compartment or cellular surface. (1). In the former case, it is suspected that, following an immunogenic event such as tumor development or infection, autoreactive CD8+ T cells from the periphery drive disease pathology through the infiltration of the central nervous system (CNS) and direct recognition of antigens presented by HLA class I molecules on neurons(2–4). In this context, tumor- or pathogen-related antigens would activate peripheral CD8+ T cells, neurological damage would arise without the necessity of an initial CNS-related insult (3). As compared to AIE with surface-targeting antibodies, AIE with intracellular-targeting antibodies show a higher brain infiltration of cytotoxic CD8+ T cells with a higher number of granzyme B+ T cells affixed to neurons and a correlation of those CD8+ T cells with neuronal damage (2, 3, 5). Oligoclonal CD8+ T cell expansion has been reported in the blood, brain tissue, and dorsal root ganglia of patients with AIE (6–8). Similarly, other paradigmatic neuroimmunological diseases such as Rasmussen encephalitis (RE) and Susac’s syndrome (SuS) also display oligoclonal CD8+ T cell expansion thus suggesting a common CD8+ T cell-driven pathology in these disorders (9, 10).

While the phenotype and T cell repertoire of CD8+ T cells in RE and SuS have been vastly studied (9, 10), similar studies with in-depth CD8+ T cell assessment are mostly lacking for AIE. Despite many efforts, neuron-reactive CD8+ T cells have not been consistently found in AIE patients. Circulating cdr2 (Yo)- and HuD-specific CD8+ T cells have been identified in paraneoplastic cerebellar degeneration (PCD) and anti-Hu encephalitis patients, respectively (6, 11–13). Yet, other groups have also contested evidence for circulating CD8+ T cells against both these antigens (14–16).

In summary, evidence regarding the phenotype of allegedly autoreactive pathogenic CD8+ T cells in AIE is scarce and identification of neuron-reactive CD8+ T cells is still elusive (6–10). One of the reasons for the limited results published so far is that previous studies were restricted to testing CD8+ T cell reactivity against the same antigens as the ones recognized by antibodies. Yet, there is no evidence that both the humoral and the cellular immune response should necessarily be directed against the very same protein. Additionally, whilst certain authors assessed a broad range of candidate epitopes by generating peptide libraries and transfecting antigen-presenting cells (APCs) with HLA molecules, these methods have never been applied to studying AIE and also have limitations (17). In particular, these techniques require the pre-selection of a restricted number of antigens and HLA alleles. Furthermore, these techniques coupled with subsequent conventional readouts usually are not able to detect CD8+ T cell clonotypes that are present at a frequency lower than 0.01% (18).

Given the state of the art, we believe that there is a need to develop novel methodologies to address the limitations of current techniques.. First, the search for neuron-reactive CD8+ T cells must be conducted in a fully human and autologous setting; second, the experimental setup must include the largest possible range of neuronal antigens to develop an unbiased approach. Third, this approach should be able to take into account protein isoforms or post-translational modifications that can affect antigen recognition by CD8+ T cells (19, 20). To fulfill all these criteria, we developed a method to assess CD8+ T cell activation against a broad repertoire of antigens naturally produced and presented by autologous neurons.

## Results

To assess the global CD8+ T cell response against neurons, we resort to the use of autologous human induced pluripotent stem cell (hiPSC)-derived neurons (21, 22) as APCs. First, to validate this approach, we generated hiPSCs from six healthy donors (HD) and differentiated these hiPSCs into neurons. We then assessed their capacity to trigger antigen-dependent CD8+ T cell activation. Upon stimulation with IFN-γ and TNFα, we found that hiPSC-derived neurons display HLA class I molecules at their surface (Figure 1a and 1b). We next assessed the capacity of hiPSC-derived neurons to activate CD8+ T cells in an antigen-dependent manner. For this, we performed an overnight stimulation of *ex vivo* CD8+ T cells with autologous neurons pulsed with CD8+ T cell-restricted viral peptide pools (see Supplementary Table 1). In this assay, we were able to elicit an IFN-γ secretion by CD8+ T cells that correlates with the secretion induced in total PBMCs during a standard IFN-γ ELISpot assay, demonstrating the capacity of neurons to trigger antigen-dependent activation of CD8+ T cells (Figure 1c).

**Figure 1.**
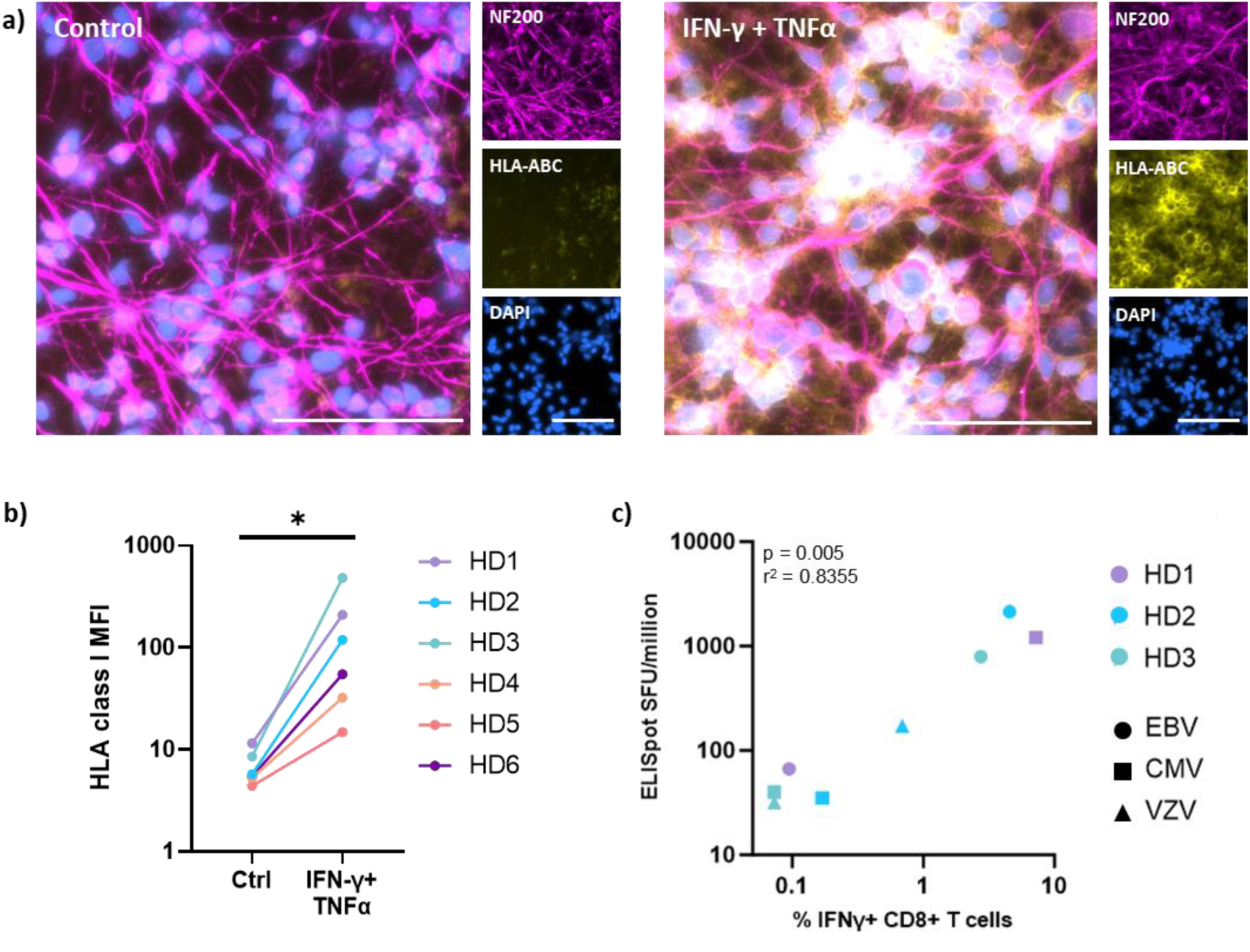
Human iPSC-derived neurons trigger antigen-dependent CD8+ T cell activation. a) Representative fluorescence microscopy images of hiPSC-derived neurons from HD6 immunostained for NF-200 (magenta), HLA-ABC (yellow) and DAPI (blue), untreated (control) or treated with IFN-γ and TNFα for 48 hours. Bar represents 100μm. b) Flow cytometry analysis of HLA class I expression (antibody against HLA-A, B and C, (clone W6/32)) of hiPSC-derived neurons from 6 healthy donors (HD1-HD6) untreated (control) or treated with IFN-γ and TNF-α for 48 hours (Wilcoxon matched-pairs test, p value = 0,0156). c) Correlation of the number of spot-forming units (SFU)/million PBMCs in an IFN-γ ELISpot assay (y axis) versus the percentage of IFNγ+ CD8+ T cells (x axis). PBMCs and neurons-T cell cocultures were pulsed overnight with a pool of CD8+ T cell-restricted immunodominant viral epitopes from EBV, CMV and VZV. Correlation was done running a Pearson correlation test.

As neuron-reactive CD8+ T cell clonotypes are also present in healthy subjects (23), we initially sought to demonstrate the feasibility of our method in a cohort of HD. To identify neuron-reactive CD8+ T cell clonotypes, we developed a coculture system between PBMCs and autologous hiPSC-derived neurons and monitored CD8+ T cell proliferation after *in vitro* autologous coculture. We assessed the TCR-β chain repertoire of CD8+ T cells after 14 days of culture and compared it to the *ex vivo* CD8+ T cell repertoire. We identified CD8+ T cell clonotypes that were expanded in the presence of autologous neurons (Figure 2a). We first performed this assay using PBMCs from 6 healthy donors and performed conventional TCR-β chain repertoire metric assessments (Shannon entropy: a measure of TCR repertoire diversity; clonality: a measure of evenness of TCR chain distribution). We observed a global yet non-significant decrease of the Shannon entropy (average of 11.095 *ex vivo* vs 9.96 at 14 days, p-value: 0.21) while the clonality remained stable (average of 0.3 ex vivo vs 0.25 at 14 days, p-value: 0.25) in the TCR repertoire after the culture (Figure 2b).

**Figure 2.**
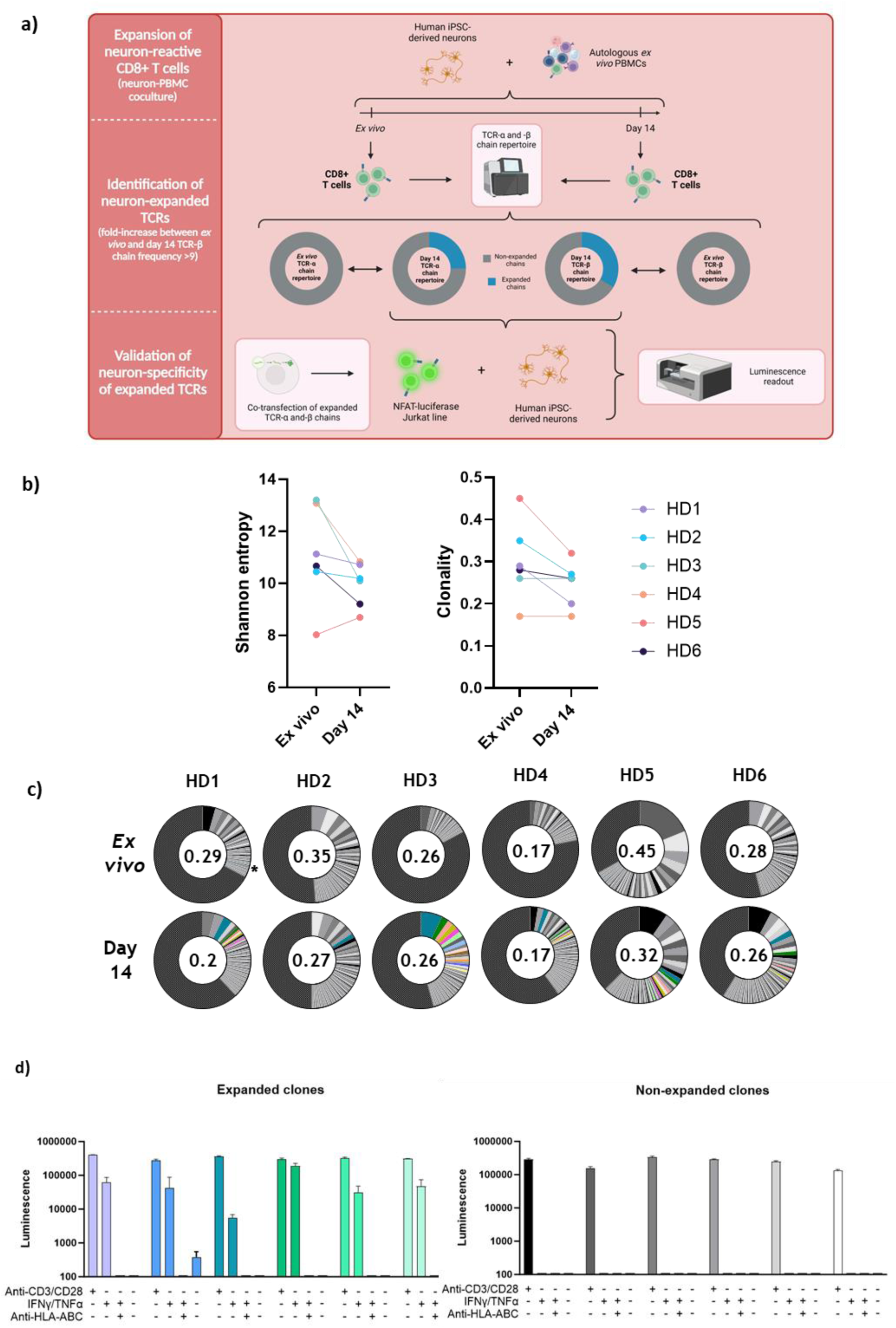
Human iPSC-derived neurons trigger the expansion of rare neuron-reactive CD8+ T cell clonotypes in healthy donors. a) Schematic representation of the experimental workflow for the identification of neuron-reactive CD8+ T cell clonotypes. First, coculture of PBMC with autologous neurons is performed for 14 days. Second, TCR–β repertoires at day 14 of coculture are compared to *ex vivo* TCR–β repertoires to identify expanded TCR-β chains. Third, paired TCR-α chains of expanded clonotypes are identified by scTCRseq (not shown) and both TCR-α and TCR–β chains are co-transfected into NFAT-luciferase reporter Jurkat CD8+ T cells which are then cocultured overnight with neurons. Luminescence values are then measured on a multimode microplate reader (see methods section for details). b) Shannon entropy (left) and repertoire clonality (right) of TCR-β chains from CD8+ T cells *ex vivo* and after 14-day co-culture between PBMCs and autologous hiPSC-derived neurons for all 6 healthy donors (HDs). Significant differences were tested with a Wilcoxon matched-pairs test, none resulted significant. c) TCR-β chain repertoire analysis of the CD8+ T cells from 6 HD *ex vivo and* after 14-day coculture. Each slice represents a unique TCR-β sequence, and the size of the slice represents the frequency of each sequence in comparison with total TCR-β sequences. Colored slices represent TCR-β chains that underwent a 9-fold expansion as compared to *ex vivo* frequencies, and that constitute >0.5% of total TCR-β after 14-day coculture. *Ex vivo* TCR-β sequences are represented with the same colors as corresponding expanded TCR-β sequences. The asterisk indicates the only TCR-β representing more than 0.5% *ex vivo*. The value at the center of each pie chart represents the clonality of each TCR-β repertoire. All TCR-β sequences <1:1000 are grouped into one slice (black and grey patterned slice). d) Luminescence values of TCR-transfected NFAT-luciferase Jurkat cells after overnight coculture with autologous neurons. Left panel: overnight culture with Jurkat bearing TCRs that were identified as expanded in HD4. Right panel: overnight culture with Jurkat bearing TCRs with prevalent TCR-β chain frequencies at day 14 yet non-expanded in HD4. Each color/grey shading represents an individual TCR. Experimental conditions include a transactivating anti-CD3/CD28 antibody without neurons, neurons treated and non-treated with IFN-y/TNFα (thus HLA class I positive and negative respectively or with addition of a blocking anti-HLA ABC antibody (clone W6/32).

We then analyzed the TCR β-chain sequences to identify specific CD8+ T cell clonotypes that would have been expanded upon coculture with autologous neurons. We considered a clonotype to be expanded *in vitro* if TCR-β chain frequency presented with a ≥9-fold expansion as compared to *ex vivo* samples. After coculture, we observed several TCR-β chains whose frequencies were significantly increased after 14 days of culture (Figure 2c). Focusing on TCR-β chains representing >0.5% of total repertoire at day 14, we could highlight a specific enrichment with a mean of 5.33 clonotypes per donor (range 1 to 12), representing from 0.5% to 7.2% of the total repertoire (Figure 2c). Only one TCR β-chain was found at an *ex vivo* frequency >0.01% (HD1 – TCR-β chain: 0.13%, very fine blue band, Figure 2C). To ensure that these expanded clonotypes recognize neurons, we restimulated CD8+ T cells with autologous neurons and sorted IFN-γ+ CD8+ T cells in 96-well plates at one cell per well. We next amplified the mRNA sequences encoding for the TCR-α and TCR-β chains in each well separately in order to reconstruct the TCRs of these cells. Using this approach, we could selected 12 TCRs from HD4 clonotype, which we cloned into Jurkat cells to assess their reactivity. We selected 6 clonotypes that we thought would recognize neurons based on their expansion kinetic *in vitro* while the 6 other clonotypes were classified as non-expanded (did not display an exponential increase between day 7 and day 14 of coculture, data not shown) (Figure 2e). When cultured with autologous neurons, 6/6 TCRs extracted from neuron-expanded CD8+ T cells displayed a specific activation. The addition of an anti-HLA class I blocking antibody, preventing the TCR-HLA interaction specifically abrogated this activation. Similarly, neurons that were not stimulated with IFN-γ and TNFα, thus not presenting antigens through HLA class I molecules, did not elicit any TCR activation (Figure 2d). Conversely, TCRs cloned from non-expanded (i.e. fold-increase <2 compared to *ex vivo*) CD8+ T cells at day 14 did not activate in contact with autologous neurons thus demonstrating that our technique specifically expands neuron-reactive CD8+ T cells (Figure 2d, right panel).

Next, we looked for the presence of neuron-reactive CD8+ T cells in a patient suffering from anti-Ri AIE (Ri01), in which the autoantibodies target an intracellular antigen. As described for HD, we generated hiPSC-derived neurons that upregulated HLA class I molecules upon IFN-γ and TNFα exposure (Figure 3a). We next performed our PBMC-neuron coculture assay to assess whether we could identify neuron-reactive CD8+ T cells in this patient. We performed the assay using PBMCs drawn at two time-points: i) at the time of disease (Ri-dis), and ii) at remission (5 months later, when the patient was treated by mycophenolate mofetil and corticosteroids (Ri-rem)). We found no significant difference in the *ex vivo* Shannon entropy or clonality of the TCR-β chain repertoire as compared to HDs. Yet, in Ri-dis the Shannon entropy decreased from 7.45 *ex vivo* to 4.47 at day 14 and clonality of the Ri-dis sample strikingly increased by 1.73-fold (0.34 *ex vivo* and 0.6 at day 14). In Ri-rem, Shannon entropy (6.13 *ex vivo* and 6.72 at day 14) and clonality (0.46 *ex vivo* and 0.42 at day 14) remained stable. This was also the case for the six HDs where TCR-β chain metrics either decreased or remained stable (Figure 3b). In sharp contrast with HDs, CD8+ T cells from the Ri-dis timepoint revealed the monoclonal expansion of one TCR-β chain, representing 58.81% of the total repertoire at day 14 (vs 6.47% *ex vivo*) (clonotype Figure 3c (in blue)). Interestingly, even if this chain (blue striped band) was present at a high frequency *ex vivo* at both timepoints, it did not expand when autologous neurons were cocultured with PBMCs from Ri-rem timepoint (12.16% at day 14 vs 15.56% *ex vivo*, Figure 3c). However, CD8+ T cells taken at the time of Ri-rem displayed an oligoclonal expansion of 6 other TCR-β chains (Figure 3c). These data suggest that a specific functional impairment of the major clonotype expanded at the disease peak occurred under immunosuppressive treatment.

**Figure 3.**
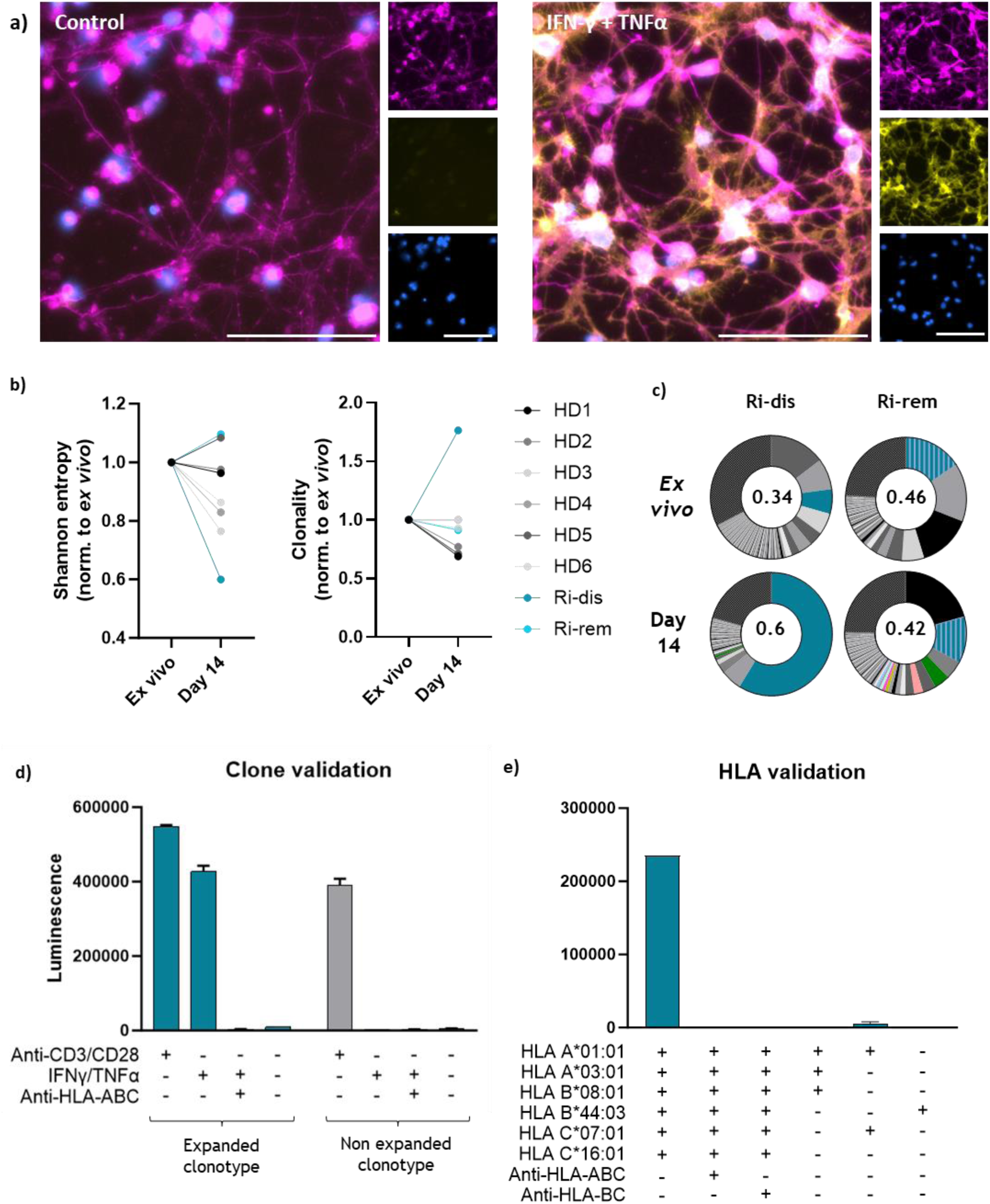
Neuron-reactive CD8+ T cells are clonally expanded in anti-Ri autoimmune encephalitis *ex vivo* and after coculture with autologous hiPSC-derived neurons. a) Representative fluorescence microscopy images of hiPSC-derived neurons from Ri01 immunostained for NF-200 (magenta), HLA-AB C (yellow) and DAPI (blue) untreated (control) or treated with IFN-γ and TNFα for 48 hours. Bar represents 100μm. b) Shannon entropy (left) and repertoire clonality (right) of TCR-β chains from CD8+ T cells *ex vivo* and after 14-day co-culture. Here, these measures are represented as relative changes as compared to *ex vivo.* c) TCR-β chain repertoire analysis of the CD8+ T cells from Ri01 patient (two timepoints: at disease (Ri-dis) and at remission (Ri-rem)), *ex vivo* and after 14-day of co-culture. Each slice represents a unique TCR-β sequence, and the size of the slice represents the frequency of each sequence in comparison with total TCR-β sequences. Colored slices represent TCR-β chains that underwent >9-fold expansion compared to *ex vivo* frequencies, and that constitute >0.5% of total TCR-β after 14-day co-culture. *Ex vivo* TCR-β sequences are represented with the same colors as corresponding expanded TCR-β sequences. The value at the center of each pie chart represents the clonality of each TCR-β repertoire. All TCR-β sequences <1:1000 are grouped into one slice (black and grey patterned slice). d) and e) Luminescence values of TCR-transfected NFAT-luciferase Jurkat cells after overnight coculture with neurons. Experimental conditions include a transactivating anti-CD3/CD28 antibody without neurons, neurons treated and non-treated with IFN-y/TNFα (thus HLA class I positive and negative respectively) or with addition of a blocking anti-HLA ABC antibody (clone W6/32). d) Left: expanded TCR; Right: most frequent yet non-expanded TCR at day 14. e) Luminescence values of NFAT-luciferase Jurkat cells transfected with the neuron-reactive TCR after coculture with autologous or partially HLA-matched neurons from 3 different healthy donors (HD2: HLA-A*01:01+, A*03:01+, B*08:01+; HD4: HLA-A*01:01+, C*07:01+; HD6: HLA-B*44:03+). In some conditions anti-HLA-ABC (clone W6/32) or anti-BC (clone B1.23.2) blocking antibodies were added.

To verify if the expanded TCR β-chain belonged to a TCR specific for neurons, and since one concomitant TCR-α chain was also expanded at day 14, we cloned the paired TCR α- and β-chains into Jurkat cells and assessed their activation. Upon overnight coculture-with neurons from Ri01, we could validate that this TCR recognizes neurons and that the antigen-dependent activation was blocked by the addition of anti-HLA class I antibody (Figure 3d). To explore which HLA allele mediated this recognition, we performed the overnight coculture with autologous neurons and added either a HLA-A, -B and -C blocking antibody, or just a HLA-B and -C blocking one. Both antibodies abolished TCR activation, highlighting that the recognition of neurons was restricted either by HLA-B or HLA-C (Figure 3d). To narrow down on HLA restriction, we conducted a coculture assay using partially HLA-matched neurons obtained from HDs (HLA A*01:01; A*03:01; B*08:01; B*44:03 and C*07:01). Since all HLA alleles from the AIE patient were covered except one (HLA-C*16:01), and since we could not observe any TCR activation, we concluded that this clonotype recognized a neuronal antigen through binding to HLA-C*16:01.

Next, to characterize the phenotype of this clonotype, we performed single cell RNA sequencing (scRNAseq) on *ex vivo* CD8+ T cells from Ri01-dis and Ri01-rem. Unbiased clustering at both timepoints revealed 14 distinct clusters (Figure 4a). Subsequently, we sought to annotate these clusters based on gene expression levels of specific markers conventionally used to classify CD8+ T cell subsets including antigen-primed (KLRG1, IL2RB), effector (PRF1, GZMB, CXCR3, FAS), naive (CCR7, SELL, BCL2) or memory (CCR7, CD27, TCF7) CD8+ T cells (Figure 4b). We could identify the neuron-reactive T cell clonotype as the 8th most frequent clonotype among all CD8+ T cells that were sequenced (=P1-C8, Figure 4d). To gain insight into the phenotype of clonotype P1-C8, we compared its gene expression profile to the one of the top 10 most frequent clonotypes . Interestingly, these clonotypes were mainly grouped within the same clusters (predominantly clusters 1, 3, 5,6 and 7, Figure 4c and 4d), which all displayed robust expression of genes associated with effector cytotoxic T cells (*KLRG1, PRF1, GZMB*) and weak expression of naïve (*CCR7, SELL, BCL2*) and central memory (*CCR7, CD27, TCF7*) T cell markers (Figure 4c).

**Figure 4.**
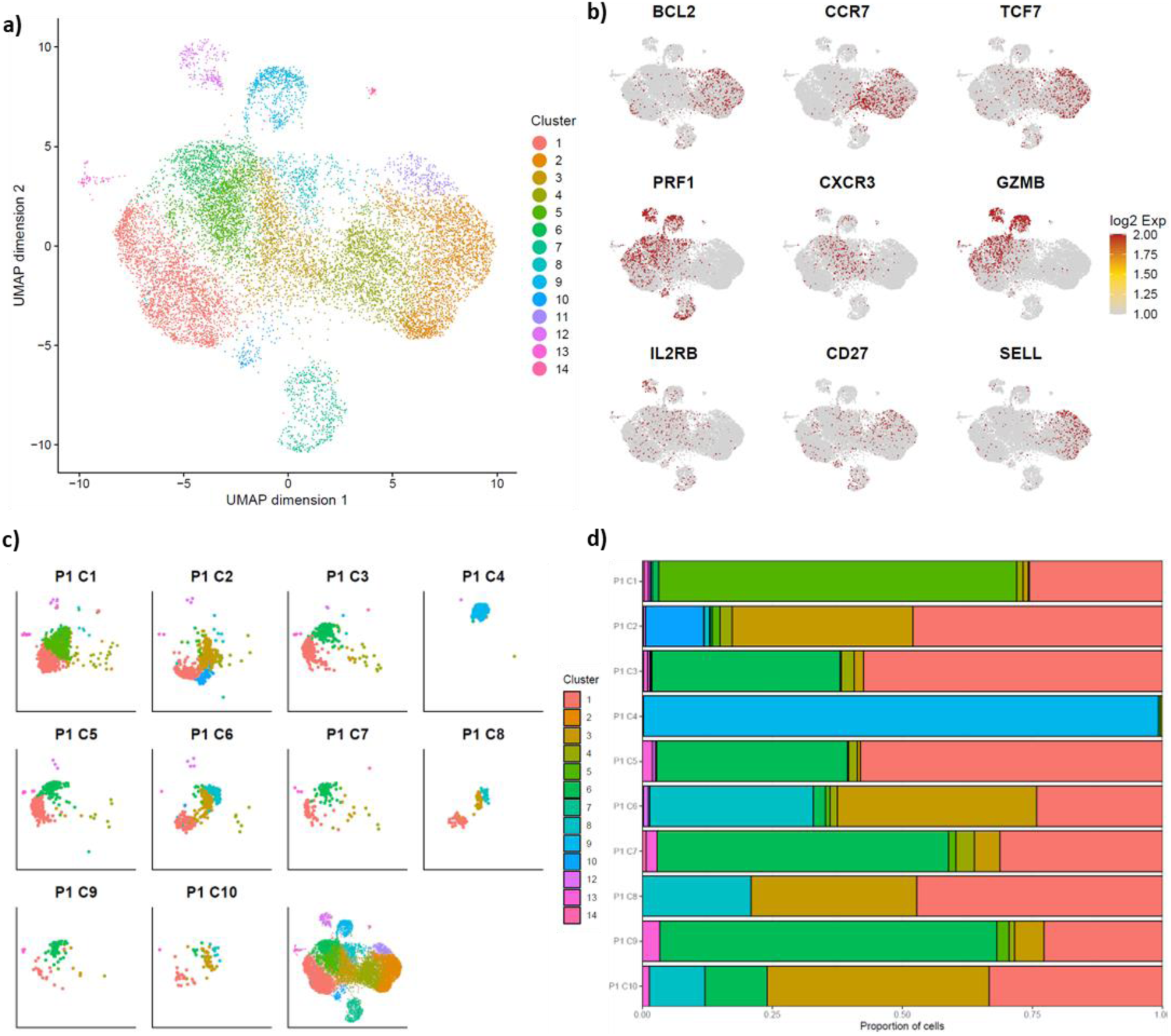
Neuron-reactive CD8+ T cells present an effector cytotoxic phenotype in anti-Ri autoimmune encephalitis. Single-cell RNA sequencing analysis of *ex vivo* CD8+ T cells from Ri01 at disease timepoint. a) UMAP with unsupervised clustering of total CD8+ T cells resulting in 14 different clusters. b) Distribution on UMAP of gene expression levels of individual markers conventionally used to classify either effector cytotoxic (PRF1, GZMB), naïve (CCR7, SELL, BCL2), activated (CXCR3, IL2RB) or central (CCR7, CD27, TCF7) CD8+ T cells. c) Distribution of the top 10 most frequent clonotypes on the UMAP projection colored by cluster. Clonotype P1-C8 represents identified neuron-reactive T cell clonotype. d) Distribution of the top 10 most frequent clonotypes *ex vivo* across all 14 clusters.

Having confirmed that the P1-C8 clonotype displayed a cytotoxic phenotype as it would be expected for pathogenic CD8+ T cells in AIE, we went on to characterize the differences between this specific clonotype and the rest of the cytotoxic most frequent clonotypes. Statistical analyses revealed that the P1-C8 clonotype displayed a unique profile of dysregulated genes discriminating it from the other 10 most frequent clonotypes (Figure 5a). Of note, P1-C2 clustered the closest to P1-C8, yet this clonotype was not expanded after coculture with neurons and did not recognize neurons after TCR cloning in Jurkat cells and overnight stimulation (Figure 3d). Detailed gene analysis highlighted an enrichment in transcription factors (*IFRD1, RUNX3, ELF1, MIF, EOMES, TOX, IRF1, KLF2*), interferon signaling-related genes (*IRF1, IFRD1, IFGNRA*), activation markers (*KLRG1, KLRB1, CD74, AKNA, KLRF1*) as well as the natural killer cytotoxicity marker *KIR3DL1* (Figure 5b). Strikingly, the combination of elevated *KIR3DL1 and TOX* expression, together with *KLRG1* and decreased expression of *IL7RA* and *ANXA1* provided with a unique signature of the P1-C8 clonotype (Figure 5c). Thus, our data highlight that the one clonotype that recognizes neurons in this patient harbors a strongly activated and cytotoxic phenotype coupled with a specific differentiation program as demonstrated by the unique combination of expressed genes. Additionally, the increased expression of key regulators of the interferon signaling pathway suggests that this clonotype may be particularly primed by IFNγ.

**Figure 5.**
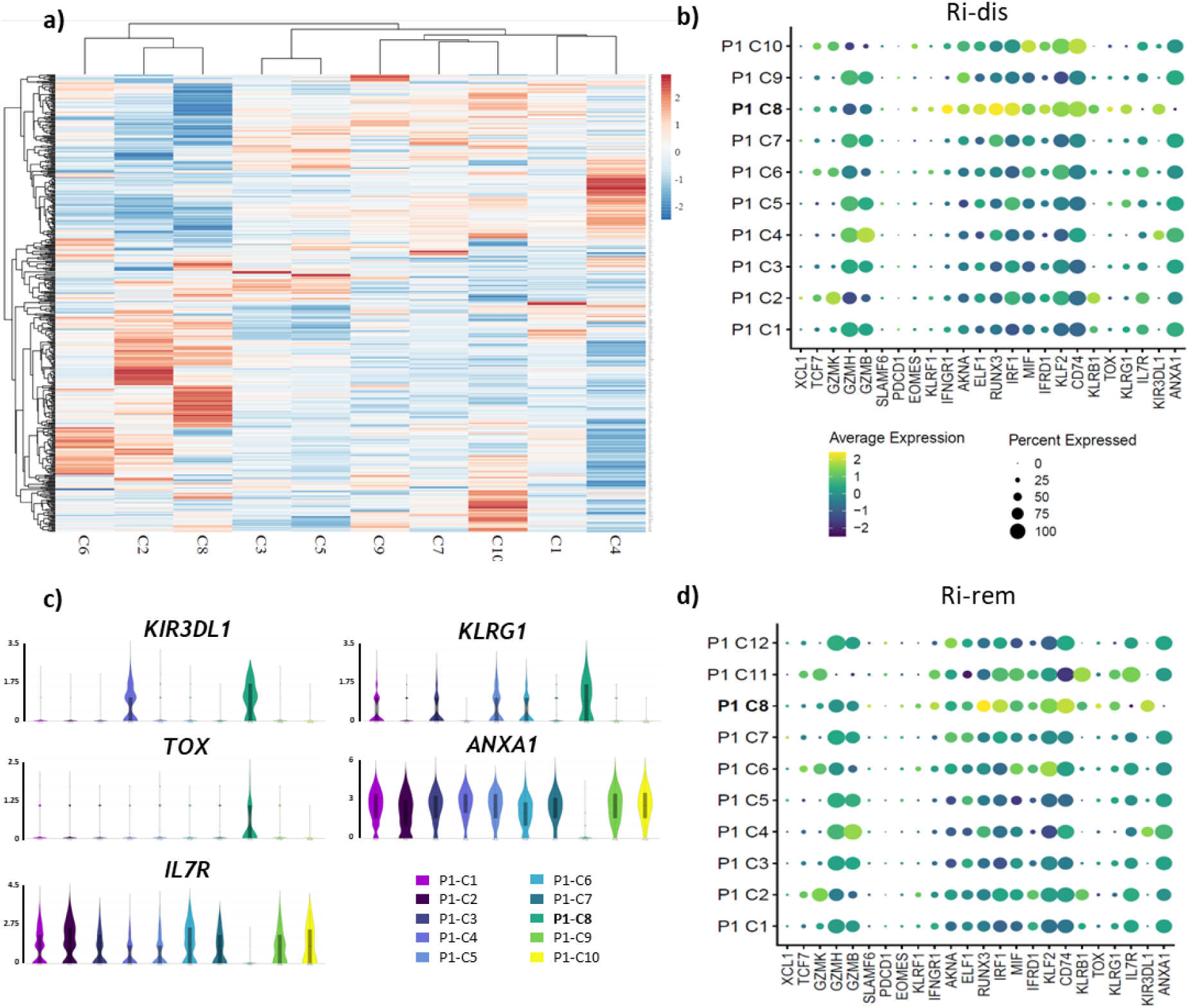
The neuron-reactive CD8+ T cell clonotype displays a unique encephalitogenic signature that persists after disease remission. Comparative single-cell RNA sequencing analysis of most frequent *ex vivo* CD8+ T cells clonotypes from Ri-dis and Ri-rem a) Heatmap of all dysregluated genes (logFC>2, adjusted p value<0.05) among the top 10 most prevalent clonotypes for Ri-dis. b) Dotplot displaying standardized average log2 gene expression per clonotype at the Ri-dis timepoint for selected genes (significantly dysregulated or not). Dot color represents the average gene expression among the cells of each clonotype, from dark blue (least expressed) to yellow (most expressed). Dot size represents the percentage of cells expressing each gene. c) Violin plots representing gene expression levels of the five most discriminant genes KIR3DL1, KLRG1, TOX (upregulated) and ANXA1, IL7R (downregulated) between P1-C8 and the other top 9 clonotypes. Gene expression is plotted as log2 gene expression. Clonotypes are represented from left to right based on decreasing order of *ex vivo* prevalence. d) Same as b) but analysis performed on Ri-rem timepoint.

Given the gap of knowledge regarding the evolution of the immune response in AIE after the peak of disease, we also performed scRNAseq on *ex vivo* CD8+ T cells of Ri-rem. Interestingly, we did not identify any difference in the global profile of CD8+ T cells. Nine out of the 10 most frequent clonotypes present at Ri-dis were still present in Ri-rem and were still mostly regrouped in the same clusters. When looking at the signature of the P1-C8 clonotype, we observed a similar profile, still harboring the P1-C8 signature previously identified, however transcription factors including *EOMES, ELF1, IFRD1, MIF and KLF2* were downregulated (Figure 5d). These data uncovered the persistence of the P1-C8 pathogenic clonotype with limited changes in its phenotype over time.

Finally, we wondered whether these findings could be extended to other patients suffering from anti-Ri AIE. We thus profiled the CD8+ T cells from another patient (Ri02) with Ri-AIE by scRNAseq and conducted the same analysis as for the previous patient. We identified 11 distinct clusters (Figure 6a and 6). Again, the 10 most frequent clonotypes were regrouped in the same clusters displaying an activated cytotoxic phenotype (i.e. *KLRG1, PRF1, GZMB*) (data not shown). Next, we sought to explore whether we could find one or several clonotypes displaying the same signature as the P1-C8 clonotype from Ri01. Strikingly, clonotype P2-C5 displayed the exact same profile with increased expression of *KIR3DL1, KLRG1, TOX* and decreased *IL7RA* and *ANXA1* (Figure 6b). In addition, P2-C5 clonotype also showed upregulation of several markers upregulated by the P1-C8 clonotype including CD74 and IRF1, Figure 6c).

**Figure 6.**
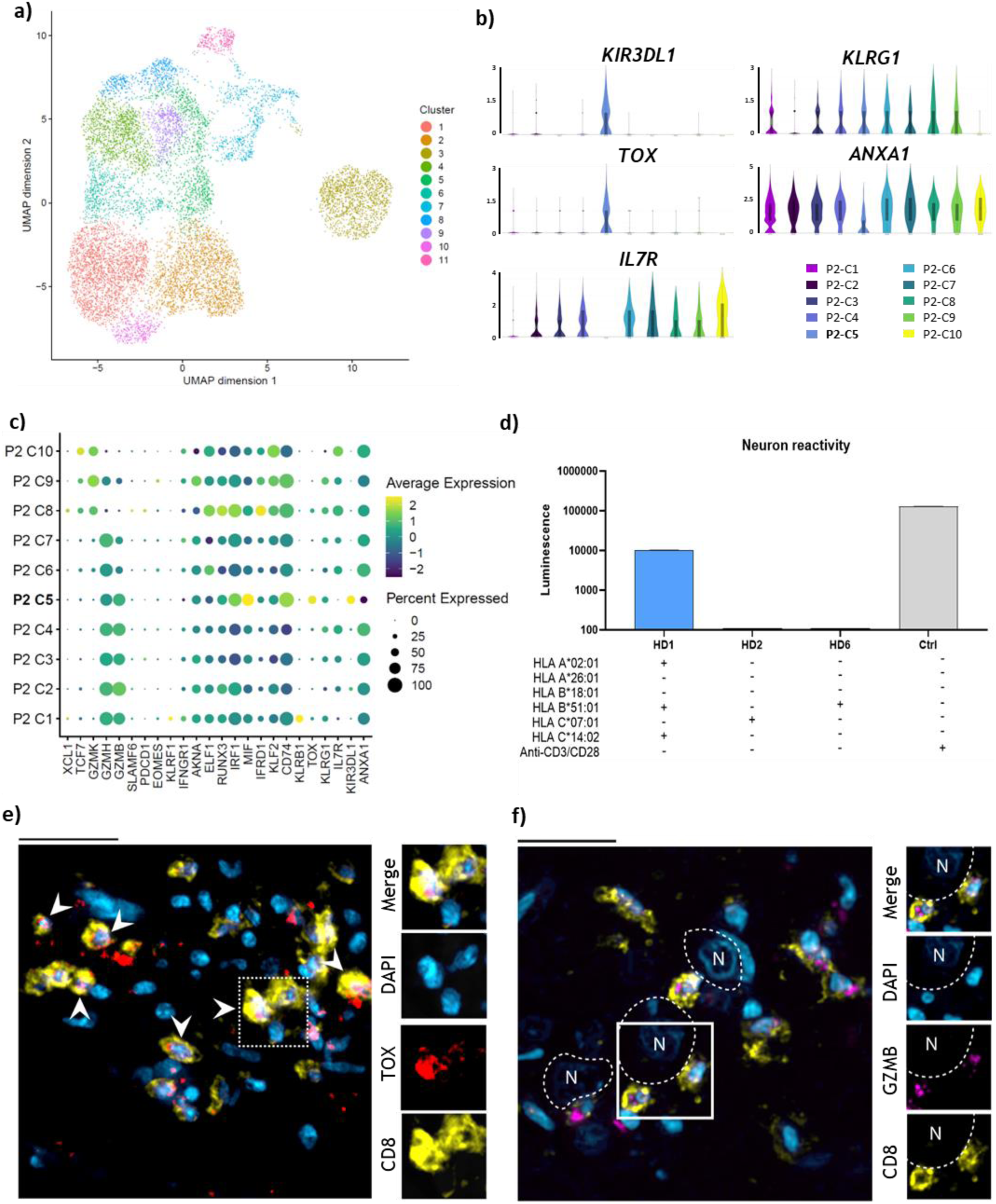
TOX+ CD8+ T cells recognize neurons and infiltrate the brain of a second patient with anti-Ri autoimmune encephalitis. a-c) Single-cell RNA sequencing analysis of *ex vivo* CD8+ T cells from Ri02. a) Unsupervised clustering of total CD8+ T cells resulting in 11 different clusters. b) Violin plots representing gene expression levels of the five most discriminant genes identifying P1-C8 (Ri01) and applied to the top 10 clonotypes found *ex vivo* in patient Ri02. Clonotypes are represented from left to right based on decreasing order of *ex vivo* prevalence. Gene expression is plotted as log2 gene expression. c) Dotplot displaying the expression of selected genes expressed by the top 10 clonotypes from Ri02. Dot color represents the average gene expression among the cells of each clonotype, from dark blue (least expressed) to yellow (most expressed). Dot size represents the percentage of cells expressing each gene. d) Luminescence values of TCR-transfected NFAT-luciferase Jurkat cells after overnight coculture with partially HLA-matched neurons from 3 different healthy donors (HD1: HLA-A*02:01 +, B*51:01+, C*14:02+; HD2: HLA-C*07:01+; HD6: HLA-B*18:01+). Experimental conditions include neurons treated with IFN-γ and TNFα (HLA class I positive – colored bars) and a transactivating anti-CD3/CD28 antibody without neurons (grey bar). e-f) Representative immunostaining of a brain lesion in autopsy tissue from Ri02. e) Brain tissue was stained for TOX (red), CD8 (yellow) and nuclei (blue). Bar length represents 20μm. f) Brain tissue was stained for GZMB (magenta), CD8 (yellow) and nuclei (blue). Bar length represents 25μm.

To assess whether P2-C5 was also neuron-reactive, we cloned this TCR into Jurkat cells and performed a coculture with partially HLA-matched neurons from HDs. Of the highest interest, we could confirm specific TCR activation against neurons expressing HLA-A*02:01 and HLA-C*14:02 (Figure 6d). Last, given the importance of TOX in triggering an encephalitogenic program in CD8+ T cells (24), we looked at whether we could find infiltrating TOX+ CD8+ T cells in the lesions of Ri02 on autopsy tissue. Strikingly, we could confirm the presence of infiltrating TOX+ CD8+ T cells thus suggesting the pathogenic role of the clonotype P2-C5. Additionally, CD8+ T cells present in the brain tissue were also GZMB+ and affixed to neurons(Figure 6f). This last set of experiments demonstrates that preponderant neuron-reactive CD8+ T cells with a cytotoxic encephalitogenic phenotype are most likely common findings in Ri-AIE and establish the role of CD8+ T cells in the pathogenesis of these disorders.

## Discussion

Here, we report on an unbiased novel method to identify neuron-autoreactive CD8+ T cells in a human-based and fully autologous system. The antigen-presenting cells used in this technique are the cells naturally producing the antigens in the body and, therefore, can present the broadest range of antigens with the right cell-type associated post-translational modifications and splicing. Indeed, such antigen modifications have been demonstrated to strongly influence T cell antigenic recognition in cancer (26), and it has been known for a long time that citrullination, for example, plays a role in antigen recognition in autoimmune diseases (26). In addition, by using fully autologous neurons, we can simultaneously screen for T cell recognition against all relevant HLA alleles for the studied cohort, including less studied HLA-C alleles.

Applying this method to screen first the presence of neuron-reactive T cells in HD, we could uncover that six out of six HD demonstrated expansion of specific T cell clonotypes upon coculture with neurons. Further validation of individual TCRs showed that expanded T cells in these conditions are indeed neuron-reactive. Although it has been previously shown that certain HDs may harbor hypocretin neuron, astrocyte or oligodendrocyte-reactive CD8+ T cells (23, 27, 28), these results highlight that the presence of circulating neuron-reactive CD8+ T cells is a highly common feature of the human TCR repertoire. Our data demonstrate that they are most strikingly present at very low frequencies in all donors studied (below 0.01%, Figure 2c). Of note, identification of these neuron-reactive T cells was possible only thanks to the very high sensitivity of our assay which depends only on the number of T cells used as input (here 10^5^ CD8+ T cells were sent for ex vivo TCR-β chain repertoire sequencing) and demonstrated sensitivity superior to 0.001%. In comparison, conventional methods are mostly limited to frequencies above 0.01% (18). As such, the methodology described here holds great promise to study the CD8+ T cell repertoire in patients with autoimmune diseases but also in HD thus providing the scientific community with the right tools to address fundamental questions on the development of autoreactive CD8+ T cells and how they can go rogue in autoimmunity.

In the second part of this study, we focused on the analysis of neuron-reactive CD8+ T cells in anti-Ri-AIE, which has never been addressed so far. Our pipeline allowed us to identify two clonotypes of neuron-reactive T cells in two different patients. Strikingly, these two clonotypes share similar features and display a phenotype indicating a pathogenic role in the disease. Both clonotypes were readily expanded *ex vivo* and belonged to the 10 most frequent clonotypes in each patient (Figures 4 and 6) contrasting with our findings in HD in whom neuron-reactive T cells were rare (frequency <0.01%, Figure 2). Such low frequencies in HDs suggest a naive phenotype (29) given that they are not expanded while our data confirm that the expanded clonotypes in Ri-AIE are effector cytotoxic CD8+ T cells. Of note, Ri-AIE being a rare occurrence, we could only include two patients in our study including Ri02 from whom we had brain material, a unique asset. The fact that we identified highly activated neuron-reactive CD8+ T cells using two different approaches reinforces the validity of our findings for Ri-AIE overall. Indeed, our coculture assay with autologous neurons allowed us to identify the P1-C8 clonotype in Ri01. Assessment of the specific gene expression of this clonotype allowed us to identify a unique cytotoxic signature, which we were able to identify in a second patient with the same disease (Ri02). The success of identifying the P2-C5 clonotype in Ri02 using the signature from Ri01 demonstrates the strong association between this set of genes and pathology-associated CD8+ T cell clonotypes. There is thus a strong rationale to think that our findings may be extended to other patients with Ri-AIE, and could provide a set of biomarkers to identify autoreactive CD8+ T cells in Ri-AIE. To our knowledge, this is the first demonstration in AIE of a specific CD8+ T cell phenotypic signature predictive of neuron-reactive TCRs.

Of the highest interest, the signature that we identify and that segregates neuron-reactive CD8+ T cells from the non-autoreactive activated/effector T cells identified *in vitro* is a strong indicator of the encephalitogenic role of these cells. Indeed, neuron-reactive CD8+ T cell clonotypes in Ri-AIE display an effector phenotype with high expression of *PRF1, GZMB, FAS, and CXCR3* thus making them able to cause neuronal damage by direct cytotoxicity. In addition, the core signature specific to neuron-reactive CD8+ T cells is composed of three genes: *KIR3DL1, KLRG1, and TOX*. *KRLG1* and *KIR3DL1* are two receptors known to be expressed by a subgroup of activated CD8+ T cells but they are more classically associated with NK cells. Interestingly, dysregulation of both receptors has previously been associated with autoimmunity and autoreactive CD8+ T cells. In combination with IL-7R^low^, KLRG1 has been described as a marker of short-lived effector cells and normally functions as an inhibitory receptor. However, it has rarely been studied in the context of autoimmunity. Yet, the few studies existing in humans, demonstrate that KLRG1+ CD8+ T cells are present in damaged tissues such as the skin and promote autoimmunity (30). Interestingly, a particular single nucleotide polymorphism (SNP) present on the gene coding for KLRG1 has been associated with protection against multiple sclerosis thus suggesting that the gene version without this particular SNP may contribute to CNS autoimmunity (31). *KIR3DL1* belongs to the Killer-cell immunoglobulin-like receptor family and is also an inhibitory receptor. KIR+ CD8+ T cells are increased in a number of autoimmune diseases and their function in this context remains unexplored (32). Interestingly, *KIR3DL1* presents a polymorphism, called *KIR3DS1*, which acts as an activator receptor instead of an inhibitory one and promotes autoimmunity (33). Last, TOX is a transcription factor involved in T cell differentiation programs. It has been associated with exhausted CD8+ T cells as well as polyfunctional T cells (34). Most importantly, TOX is a key regulator a the encephalitogenic program in CD8+ T cells as demonstrated in experimental models of brain autoimmunity and in the brain parenchyma (24, 35). As a matter of facts, the neuron-reactive T cells we found in Ri-AIE share some markers with encephalitogenic CD8+ T cells in these models. Moreover, we also demonstrated that TOX+ CD8+ T cells are found in the lesions of the second Ri-AIE for whom we had autopsy tissue, a feature also identified by Frieser et al (35). Remarkably, TOX+ CD8+ T cells were also found in MS lesions, suggesting a role in human lesion pathogenesis (24). ANXA1 is specifically downregulated in our autoreactive clonotoypes., Whilst this gene has been poorly studied in CD8+ T cells, reduced ANXA1 in CD4+ T cells has been shown to associate with inflammation and disease severity in MS (36), suggesting that this be could a common feature in CD8+ T cells and AIE, although this remains to be demonstrated.

Further analysis revealed that both pathogenic neuron-reactive T cell clonotypes recognize neurons via HLA-C molecules in Ri-AIE. The P1-C8 clonotype most likely recognizes neurons through HLA-C*16:01 and not HLA-A or HLA-B. While most of the genetic associations identified in neurologic autoimmune disorders involve class II HLAs (37, 38), it is interesting to note that the strongest HLA association in the field is the carriage of HLA-C*07/HLA-C*06 in more than 90% of patients with Susac syndrome (10, 39) and patients with Rasmussen’s encephalitis (9). Furthermore, the analysis performed on the second patient with Ri-AIE revealed that a neuronal-reactive T cell clonotype sharing similar properties as the one from Ri01 recognizes neurons through either HLA-A*02:01 or HLA-C*14:02. Both HLA-C*16:01 and HLA-C*14:02 belong to the Bw6 serotype group and share 97.5% of homology in the amino-acid sequence. In addition, they are both rare alleles carried by 3.4% and 2.2.% of the population respectively. Altogether, these pieces of evidence suggest a strong implication of HLA-C molecules in CD8+ T cell pathology in Ri-AIE.

Taken together, our findings establish that CD8+ T cells are instrumental in Ri-AIE and that neuron-reactive clonotypes express a specific gene signature predictive of HLA-C-restricted autoimmunity with. Finally, our method has the potential to advance the understanding of pathogenic CD8+ T cells in autoimmune neurological diseases, paving the way for significant insights into this intricate field, and suggests that hiPSC-derived somatic cells could be used as antigen-presenting cells to study a large variety of other autoimmune diseases.

## Methods

### Study population

Blood draw samples from six HD and two patients with anti-Ri AIE were enrolled for this project (Ri01 had blood drawn at disease (Ri-dis) and remission (Ri-rem) timepoint (Supplementary Table 1). All donors gave their written informed consent according to regulations established by the responsible ethics committee (Project COOLIN’BRAIN, CER-VD 2018-01622). Peripheral blood mononuclear cells (PBMC) from each donor were obtained by standard Ficoll-Paque (Sigma-Aldrich) gradient centrifugation and cells were liquid nitrogen-frozen in fetal bovine serum (FBS, Biowest) and DMSO (1:10, Sigma-Aldrich). PBMC from AIE patients were taken at the moment of initial disease onset and for Ri01, also during remission as part of our longitudinal follow-up.

### Generation and characterization of human-induced pluripotent stem cell (hiPSC)-derived neurons and precursors

For all six HD and Ri01 patient, hiPSCs were reprogrammed from CD71+ cells isolated from PBMCs and passed all standard quality controls as we described it previously (1, 2). Human iPSCs were then differentiated into neural precursor cells (NPCs) (1, 3) and mature neurons were then obtained through transduction of NPCs with an NGN2 lentiviral vector as described by Ho et al (4) and used in previous publications (5, 6). Human iPSC-derived neurons were characterized by immunohistochemistry as follows. First, they were plated and differentiated in polyornithine (1:5, Sigma-Aldrich)/laminine (1:500, Sigma-Aldrich)-coated tissue culture treated 24 well plates and treated with IFN-γ (1ng/ml, Miltenyi Biotec) and TNFα (1ng/ml, R&D Systems) for 48 hours to induce HLA class I enhancement (see Human-iPSC-derived neuron preparation section and Supplementary Table 5). Neurons were washed once with 500ul of cold PBS and fixated with PBS + PFA (4:100, Electron Microscopy Sciences) for 10 min at 4°C. Wells were then washed with blocking buffer (PBS + Normal goat serum (5:100, Jackson ImmunoResearch) + 0.1% Triton X-100 (Sigma-Aldrich)). Primary antibodies including rabbit IgG anti-NF200 (1:200, Sigma Aldrich, polyclonal), chicken IgY anti-MAP2 (1:200, Abcam, polyclonal) and mouse IgG2a anti-HLA-A, -B, -C (1:500, AffinityImmuno, clone W6/32) were incubated in 250ul blocking buffer with neurons overnight at 4°C. After extensive washing steps, secondary antibodies including anti-rabbit IgG (H+L) AF546 (Invitrogen), anti-chicken IgY (H+L) AF546 (Invitrogen) and anti-mouse IgG (H+L) AF488 (Invitrogen) were added for 30 minutes at RT. Counterstaining with DAPI (1:500, Sigma-Aldrich) was also performed at this stage. Then, after additional washing steps, images were acquired using a Leica DMi8 microscope and post-processed with LAS X software.

### Human-iPSC-derived neuron preparation (HLA enhancement and peptide pulsation)

Neurons were plated in polyornithine (1:5, Sigma-Aldrich)/laminine (1:500, Sigma-Aldrich)-coated tissue culture treated 48-well plates (Corning) at a density of 100’000 cells/cm^2^. To induce HLA class I upregulation, cell medium was replaced with fresh medium supplemented with IFN-γ1b (1ng/ml, Miltenyi Biotec) and TNFα (1ng/ml, R&D Systems) for 48 hrs. Some experiments required the pulsation of CNS cells with reconstituted viral peptide pools. To this end, CNS cells were pulsated with CD8+ T cell-restricted peptide pools from either EBV, CMV or VZV (1ug/ml for each peptide pool, JPT Peptide Technologies) four hours prior to culture with CD8+ T cells. EBV and CMV CD8+ T cell-restricted peptide pools were reconstituted from respectively 29 and 45 individual immunogenic peptides (JPT Peptide Technologies) (see Supplementary Table 4 for each individual peptide sequence). VZV peptide pools were acquired from a commercially available mix of 63 individual peptides (PepMix VZV, IE63, JPT Peptide Technologies). Importantly, prior to overnight incubation with CD8+ T cells, hiPSC-derived neurons were carefully washed four times with cell medium in order to remove any residual cytokines (ie. IFN-γ or TNFα) or viral peptides.

### Neuron – *ex vivo* CD8+ T cell overnight stimulation

*Ex vivo* PBMC were thawed and rested overnight in a serum-free T cell expansion medium (SFM, Thermo Fisher Scientific). After resting, CD8+ T cells were isolated by magnetic-associated cell sorting (MACS) (see MACS section below), counted and exposed to HLA class I-enhanced neurons (+/-peptide pulsation) (see CNS cell preparation section above) at a ratio of 1:1 (i.e. 100’000 CD8+ T cells for 100’000 neurons per well). Neurons and autologous CD8+ T cells were cultured together overnight, and an IFN-y secretion assay was performed (see IFN-y secretion assay section below).

### Neuron-PBMC co-culture

Autologous PBMC were thawed and rested overnight in SFM. Rested PBMC were counted, resuspended at 4 mio cells/ml and then cultured with HLA class I-enhanced neurons (see Human-iPSC-derived neuron preparation section above) at a density of 1.2mio cells/cm^2^ with IL-2 (1000UI/ml, Miltenyi Biotec) and a CD28 agonist antibody (5ug/ml, BD Biosciences, clone CD28.2) for a total of 14 days. 300ul of fresh SFM and IL-2 (1000UI/ml) was added at day 2 and then renewed at day 5. At day 7, PBMC from the co-culture were harvested, counted and resuspended at 4 mio cells/ml and then re-cultured with fresh HLA I-enhanced neurons. 300ul of fresh SFM and IL-2 (1000UI/ml) was added at day 10 and then renewed at day 12. Bulk TCR-α and -β chain repertoire sequencing was performed directly *ex vivo* as well as at days 7 and 14 of co-culture (see Bulk and single-cell TCR-α and TCR-β chain repertoire sequencing and analysis section). In some cases, an IFN-y secretion assay was performed at day 14 of co-culture and subsequent FACS was performed to isolate CD3+ CD8+ IFN-y+ fractions (see IFN-y secretion assay, Flow cytometry and Single-cell TCR-α and TCR-β chain sequencing sections).

### Magnetic-associated cell sorting (MACS)

Isolation of untouched CD8+ T cells from total PBMCs was performed using a two-step human CD8+ T cell isolation kit (Miltenyi Biotec). All steps were performed following manufacturer instructions. After MACS, additional purity checks were performed by flow cytometry (see Flow cytometry analysis section).

### IFN-γ secretion assay

This assay was performed in two separate experiments: 1) Overnight stimulation of *ex vivo* CD8+ T cells with HLA I-enhanced neurons (+- peptide pulsation); 2) Overnight culture of CD8+ T cells isolated after 14-day of neuron-PBMC co-culture with HLA I-enhanced neurons. Additional controls included HLA-unenhanced neurons or addition of a blocking anti-HLA A, -B or -C antibody (W6/32, Affinity Immuno).

IFN-γ production of CD8+ T cells was measured by adapting a commercially available IFN-γsecretion assay kit (Miltenyi Biotec). Briefly, MACS-sorted CD8+ T cells were stained with an IFN-γ Catch Reagent prior to overnight culture with neurons. Additionally, an IFN-γ Detection Antibody (1:100) was added to neuron-CD8+ T cell coculture prior to overnight incubation. If not otherwise specified, all other steps and reagent concentrations were applied following manufacturer instructions.

### Flow cytometry

For HLA class I assessment on CNS cells, neuronswere detached using TrypLE (Gibco) and washed with phosphate buffer saline (PBS) and FBS (2:100). Cells were then stained with a pan HLA-A, -B, -C antibody (2:100, Santa Cruz Biotechnology, clone W6/32), washed again with PBS + 2% FBS and fixated in PBS + PFA (4:100, Electron Microscopy Sciences).

CD8+ T cell staining was performed the day after the overnight IFN-y secretion assay or for post-MACS purity checks. Cells in suspension were harvested and washed with phosphate buffer saline (PBS) and 2% FBS. Anti-CD8a (2:100, BD Biosciences, clone RPA-T8) and anti-CD3 (2:100, BD Biosciences, clone SK7) were then added, cells were washed again with PBS + 2% FBS and fixated in PBS + 4% PFA. Viability of all cells was assessed using an aqua fluorescent reactive amine dye (4:1000, Life Technologies). Surface marker expression was assessed using an LSR II flow cytometer (BD Biosciences).

When single cell TCR sequencing was necessary (see Single-cell TCR-α and TCR-β chain sequencingsection), after overnight IFN-y secretion assay, CD8+ T cells were sorted by fluorescence-associated cell sorting (FACS) into IFN-γ+ CD3+ CD8+ T populations at 1 cell/well. For this, CD8+ T cells were stained and processed as described above and sorting was performed using a FACSAria III cytometer (BD Biosciences). No PBS-PFA fixation steps were performed in this case.

### Bulk TCR-α and TCR-β chain repertoire sequencing and analysis

MACS-isolated CD8+ T cells were suspended in 300μl Lysis/Binding Buffer (Thermo Fisher Scientific) and bulk TCR sequencing analyses were performed as described previously (7). Briefly, mRNA was isolated and amplified by in vitro transcription. 5’ adapter was added by multiplex reverse transcription and TCRs were amplified using one primer in the adapter and one in the constant region. Libraries were sequenced on Illumina instrument and TCR sequences processed using an ad *hoc Perl* script.The Shannon Entropy was calculated as follows :

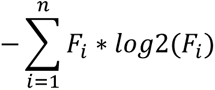

Where n is the total number of clonotype and F the clonotype frequency. Clonality, refers to 1-Pielou index, was calculated as follows :

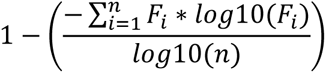

Where n is the total number of clonotypes and F the clonotype frequency.

### Single-cell TCR-α and TCR-β chain sequencing (scTCRseq)

For the pairing of the corresponding TCR-α chain, scTCRseq on CD8+ T cells at day 14 was performed. For this, IFN-y+ CD3+ CD8+ T cells were sorted by FACS at 1 cell/well into 96-well PCR plates containing 15ul of DNase/RNase-free distilled water (Invitrogen) with 0.2% Triton X-100 (Sigma-Aldrich) and an RNase inhibitor (2U/ul) (Enzymatics). For the pairing of the corresponding TCR-α chain, scTCRseq on CD8+ T cells at day 14 was performed. For this, IFN-y+ CD3+ CD8+ T cells were sorted by FACS at 1 cell/well into 96-well PCR plates containing 15 µL of DNase/RNase-free distilled water (Invitrogen) with 0.2% Triton X-100 (Sigma-Aldrich) and an RNase inhibitor (2U/µL) (Enzymatics). TCRα/β were amplified using OneStep RT-PCR kit (Qiagen) according to the manufacturer recommendation, with the following modification: a collection covering all V segments and two primers designed in alpha and beta constant regions were used for the amplification. Then, amplicons were purified using AmpureXP beads (Beckman Coulter) according to the manufacturer instruction. A second amplification was performed with the Phusion Hot Start DNA Polymerase (NEB). A forward primer, designed in the adapter sequence (added during the RT-PCR) and two reverse nested primers, designed in the constant regions, were used for the amplification. The PCR mix was composed of 1 µL of purified product, 0.4 µL of 10 mM dNTP mix (Promega), 0.4 µL of primers mix *alpha*, 0.4 µL of primers mix *beta*, 2 µL of buffer and 0.2 µL of polymerase and 5.6 primers mix H_2_O. α/β TCR chains were amplified with the following PCR cycles: 98°C for 4’, 25 x (98°C for 10’’, 55°C for 30’’, 72°C for 30’’), 72°C for 2’. PCR product was purified by adding 1 µL of ExoSAP-IT (Affymetrix) and incubating at 37°C for 15’ and then at 85°C for 15’. The third PCR was performed with the Phusion hot Start (NEB) to add the Illumina adapter and index. A mix containing 1 µL of 10 mM dNTP (Promega), 5 µL of 0.25 µM NexteraXT primer index, 3 µL of buffer, 0.2 µL of enzyme and 5.8 µL of deionized water was prepared and added directly to the purified PCR product. The second amplification was performed as follows : 98°C for 4’, 15 x (98°C for 10’’, 55°C for 30’’ 72°C for 30’’), 72°C for 2’. After purification on AmpureXP beads (Beckman Coulter), libraries were sequenced on Illumina instruments and sequences extracted using an *ad hoc* perl script. Both chains were amplified then transfected into reporter cell lines as described below (see TCR-α and -*β* chain amplification cloning and validation section).

### TCR-α and -*β* chain amplification, cloning and validation

For all HDs tested and Ri01, single TCR-α and -β chains were amplified from residual bulk RNA material as described previously (25). Briefly, two primers were designed in the CDR3 of each TCR to specifically amplify the V and the J regions. The two amplicons were combined by fusion PCR and the constant region added by a second fusion PCR.

For Ri02 sample, fully human codon-optimized DNA sequences were synthesized at GeneArt (Thermo Fisher Scientific) and served as template for *in vitro* transcription (IVT) and polyadenylation of RNA molecules as per the manufacturer’s instructions (HIScribe T7 ARCA mRNA kit (with tailing), NEB), followed by co-transfection into recipient T cells.

To validate antigen specificity and interrogate T cell reactivity, TCR-α/β pairs were cloned into recipient Jurkat cell line (TCR/CD3 Jurkat-luc cells (NFAT), Promega, in-house stably transduced with human CD8α/β and TCRα/β CRISPR-KO), as previously described (8). Jurkat cells were electroporated using the Neon electroporation system (Thermo Fisher Scientific) with the following parameters: 1,325 V, 10 ms, 3 pulses. Electroporated Jurkat cells were co-cultured with HLA-enhanced hiPSC-derived neurons at a ratio of 1:1 (i.e. 100’000 Jurkat for 100’000 neurons) in a 48 well-plate in 200ul SFM. All conditions were run in duplicates. After overnight incubation, 50’000 Jurkat cells were transferred into opaque tissue-culture treated 96-well plates (Corning) and the assay was performed using the Bio-Glo Luciferase Assay System (Promega). Additional controls include mock-(transfection with nuclease-free water) transfected Jurkat cells and co-culture with HLA-unenhanced neurons or in presence of a blocking anti-HLA ABC antibody (W6/32, AffinityImmuno). Luminescence was measured with a Multimode Microplate Reader (BioTek Synergy). As positive control for TCR activation, Jurkat cells were cultured in the presence of TransAct^TM^ (Miltenyi), i.e. anti-CD3/CD28.

### Single-cell RNA and VDJ sequencing

CD8+ T cells positively sorted by MACS (Miltenyi Biotec) were assessed for viability by AO/PI staining and counted with a Luna-FX7^TM^ Automated cell counter (Logos Biosystems).

A Chromium Next GEM Chip N (10X genomics) was loaded with the appropriate number of cells, and the sequencing libraries prepared with the Chromium Next GEM Single Cell 5’ HT Reagent Kits v2 dual index following the manufacturer’s recommendations. Briefly, an emulsion encapsulating single cells, reverse transcription reagents and cell barcoding oligonucleotides was generated. After the actual reverse transcription step, the emulsion was broken, and double stranded cDNA generated and amplified in a bulk reaction. This cDNA was fragmented, a P7 sequencing adaptor ligated, and a 5’ gene expression library generated by PCR amplification. For V(D)J sequencing, a similar approach was followed except that 2 steps of PCR based V(D)J target enrichment were performed prior to fragmentation.

Libraries were quantified by a fluorimetric method, and their quality assessed on a Fragment Analyzer (Agilent Technologies). Sequencing was performed on Illumina NovaSeq 6000 v1.5 flow cells for 28-10-10-90 cycles (read1 - index i7 - index i5 - read2) with 1% PhiX spike in. Sequencing data were demultiplexed using the bcl2fastq2 Conversion Software (v. 2.20, Illumina) and primary data analysis performed with the Cell Ranger Gene Expression pipeline (version 7.1.0, 10X Genomics). Mapping of VDJ sequences to gene expression library resulted in a total of >96.4% mapped TCR-β chains (Ri01_dis: 96.4%, Ri01_5m: 96.7% and Ri02: 96.4%) and >38.80% TCR-a chains (Ri01_dis: 53.3%, Ri01_5m: 43.2% and Ri02: 38.8%).

### CD8+ T cell clustering analysis and cell subtype annotation

Louvain graph-based clustering and differential expression analyses of the TCR-seq data were performed using 10X Genomics Loupe Browser (version v7.0.0) with default parameters. Cell type identities of clusters were deduced by manual inspection of marker gene expression . Independently, we also consolidated the VDJ from the filtered CellRanger output using scRepertoire v2.0.0 (9) (R package version 1.12.0) and filtered the gene expression data using Seurat v5.0.3 (10) to exclude cells with less than 100 or more than 6000 detected features, or more than 10% mitochondrial reads. Cells were further filtered to include only cells CD8A or CD8B expressed, or with an associated TCR sequence. We quantified clonotypes by selecting barcodes associated with the presence of a TCR-β chain and CDR3 amino acid sequence, and confirmed that the two analysis pipelines produced near identical clonotype counts). UMAP figures and violin plots were generated using Loupe Browser, bar plots and heat maps were generated using GraphPad Prism® 9 software (Version 9.1.0) and dotplots of clonotype expression were generated using Seurat v5.0.3. Code for the Seurat analyses is available at https://github.com/bdsc-tds/Perriot_Jones_2024.

### Graphical representation

Schematic representations were created with BioRender.com (Institutional license, University of Lausanne). Graphical representations of HLA class I assessments, IFN-y secretion assays and single-cell and bulk TCR sequencing were developed using GraphPad Prism® 9 software (Version 9.1.0). Regarding scRNA/VDJ sequencing, graphical representations were performed using Loupe Browser® 7 software (Version 7.0.0), with the exception of the dotplots in Figures 5 and 6, which were created using Seurat v2.0.0.

### Histology

For immunofluorescence staining, after antigen retrieval (Sodium Citrate pH6, 30 min) and blocking of unspecific binding (PBS and FBS (1:10)), PFA-fixed sections were incubated with primary antibodies (mouse IgG1 anti-CD8a (1:50, Abcam, clone C8/144B), rat anti-TOX (Thermo Fisher Scientific, clone TXRX10), mouse IgG2a anti-GZMB (1:20, Monosan, clone GrB-7). To amplify the signals of TOX, bound antibodies were visualized with appropriate species-specific Cy5-conjugated secondary antibodies for anti–rat tyramide signal amplification (TSA). Nuclei were stained with DAPI (Sigma-Aldrich) (see Supplementary Table 6). Immunostained sections were scanned using Pannoramic 250 FLASH II (3DHISTECH) Digital Slide Scanner with objective magnification of 20x. For representative images, white balance was adjusted, and contrast was linearly enhanced using the tools levels, curves, brightness, and contrast in Adobe Photoshop CC. Image processing was applied uniformly across all images within a given dataset.

## References

1. Lancaster E. The Diagnosis and Treatment of Autoimmune Encephalitis. J Clin Neurol. 2016;12(1):1–13.

2. Bien CG, Vincent A, Barnett MH, Becker AJ, Blumcke I, Graus F, et al. Immunopathology of autoantibody-associated encephalitides: clues for pathogenesis. Brain. 2012;135(Pt 5):1622–38.

3. Melzer N, Meuth SG, Wiendl H. CD8+ T cells and neuronal damage: direct and collateral mechanisms of cytotoxicity and impaired electrical excitability. FASEB J. 2009;23(11):3659–73.

4. Bernal F, Graus F, Pifarre A, Saiz A, Benyahia B, Ribalta T. Immunohistochemical analysis of anti-Hu-associated paraneoplastic encephalomyelitis. Acta Neuropathol. 2002;103(5):509–15.

5. Schwab N, Bien CG, Waschbisch A, Becker A, Vince GH, Dornmair K, et al. CD8+ T-cell clones dominate brain infiltrates in Rasmussen encephalitis and persist in the periphery. Brain. 2009;132(Pt 5):1236–46.

6. Albert ML, Darnell JC, Bender A, Francisco LM, Bhardwaj N, Darnell RB. Tumor-specific killer cells in paraneoplastic cerebellar degeneration. Nat Med. 1998;4(11):1321–4.

7. Plonquet A, Gherardi RK, Creange A, Antoine JC, Benyahia B, Grisold W, et al. Oligoclonal T-cells in blood and target tissues of patients with anti-Hu syndrome. J Neuroimmunol. 2002;122(1-2):100–5.

8. Voltz R, Dalmau J, Posner JB, Rosenfeld MR. T-cell receptor analysis in anti-Hu associated paraneoplastic encephalomyelitis. Neurology. 1998;51(4):1146–50.

9. Schneider-Hohendorf T, Mohan H, Bien CG, Breuer J, Becker A, Gorlich D, et al. CD8(+) T-cell pathogenicity in Rasmussen encephalitis elucidated by large-scale T-cell receptor sequencing. Nat Commun. 2016;7:11153.

10. Gross CC, Meyer C, Bhatia U, Yshii L, Kleffner I, Bauer J, et al. CD8(+) T cell-mediated endotheliopathy is a targetable mechanism of neuro-inflammation in Susac syndrome. Nat Commun. 2019;10(1):5779.

11. Rousseau A, Benyahia B, Dalmau J, Connan F, Guillet JG, Delattre JY, et al. T cell response to Hu-D peptides in patients with anti-Hu syndrome. J Neurooncol. 2005;71(3):231–6.

12. Plonquet A, Garcia-Pons F, Fernandez E, Philippe C, Marquet J, Rouard H, et al. Peptides derived from the onconeural HuD protein can elicit cytotoxic responses in HHD mouse and human. J Neuroimmunol. 2003;142(1-2):93–100.

13. Roberts WK, Deluca IJ, Thomas A, Fak J, Williams T, Buckley N, et al. Patients with lung cancer and paraneoplastic Hu syndrome harbor HuD-specific type 2 CD8+ T cells. J Clin Invest. 2009;119(7):2042–51.

14. de Beukelaar JW, Verjans GM, van Norden Y, Milikan JC, Kraan J, Hooijkaas H, et al. No evidence for circulating HuD-specific CD8+ T cells in patients with paraneoplastic neurological syndromes and Hu antibodies. Cancer Immunol Immunother. 2007;56(9):1501–6.

15. de Jongste AH, de Graaf MT, Martinuzzi E, van den Broek PD, Kraan J, Lamers CH, et al. Three sensitive assays do not provide evidence for circulating HuD-specific T cells in the blood of patients with paraneoplastic neurological syndromes with anti-Hu antibodies. Neuro Oncol. 2012;14(7):841–8.

16. Carpenter EL, Vance BA, Klein RS, Voloschin A, Dalmau J, Vonderheide RH. Functional analysis of CD8+ T cell responses to the onconeural self protein cdr2 in patients with paraneoplastic cerebellar degeneration. J Neuroimmunol. 2008;193(1-2):173–82.

17. Prinz JC. Immunogenic self-peptides - the great unknowns in autoimmunity: Identifying T-cell epitopes driving the autoimmune response in autoimmune diseases. Front Immunol. 2022;13:1097871.

18. Moodie Z, Price L, Gouttefangeas C, Mander A, Janetzki S, Lower M, et al. Response definition criteria for ELISPOT assays revisited. Cancer Immunol Immunother. 2010;59(10):1489–501.

19. Petersen J, Purcell AW, Rossjohn J. Post-translationally modified T cell epitopes: immune recognition and immunotherapy. J Mol Med (Berl). 2009;87(11):1045–51.

20. Sidney J, Vela JL, Friedrich D, Kolla R, von Herrath M, Wesley JD, et al. Low HLA binding of diabetes-associated CD8+ T-cell epitopes is increased by post translational modifications. BMC Immunol. 2018;19(1):12.

21. Dolmetsch R, Geschwind DH. The human brain in a dish: the promise of iPSC-derived neurons. Cell. 2011;145(6):831–4.

22. Ho SM, Hartley BJ, Tcw J, Beaumont M, Stafford K, Slesinger PA, et al. Rapid Ngn2-induction of excitatory neurons from hiPSC-derived neural progenitor cells. Methods. 2016;101:113–24.

23. Pedersen NW, Holm A, Kristensen NP, Bjerregaard AM, Bentzen AK, Marquard AM, et al. CD8(+) T cells from patients with narcolepsy and healthy controls recognize hypocretin neuron-specific antigens. Nat Commun. 2019;10(1):837.

24. Page N, Klimek B, De Roo M, Steinbach K, Soldati H, Lemeille S, et al. Expression of the DNA-Binding Factor TOX Promotes the Encephalitogenic Potential of Microbe-Induced Autoreactive CD8(+) T Cells. Immunity. 2018;48(5):937–50 e8.

25. Hanada K, Yewdell JW, Yang JC. Immune recognition of a human renal cancer antigen through post-translational protein splicing. Nature. 2004;427(6971):252-6.

26. Moon JS, Younis S, Ramadoss NS, Iyer R, Sheth K, Sharpe O, et al. Cytotoxic CD8(+) T cells target citrullinated antigens in rheumatoid arthritis. Nat Commun. 2023;14(1):319.

27. Standifer NE, Ouyang Q, Panagiotopoulos C, Verchere CB, Tan R, Greenbaum CJ, et al. Identification of Novel HLA-A*0201-restricted epitopes in recent-onset type 1 diabetic subjects and antibody-positive relatives. Diabetes. 2006;55(11):3061–7.

28. Sabatino JJ, Jr., Wilson MR, Calabresi PA, Hauser SL, Schneck JP, Zamvil SS. Anti-CD20 therapy depletes activated myelin-specific CD8(+) T cells in multiple sclerosis. Proc Natl Acad Sci U S A. 2019;116(51):25800–7.

29. Pittet MJ, Valmori D, Dunbar PR, Speiser DE, Lienard D, Lejeune F, et al. High frequencies of naive Melan-A/MART-1-specific CD8(+) T cells in a large proportion of human histocompatibility leukocyte antigen (HLA)-A2 individuals. J Exp Med. 1999;190(5):705–15.

30. Hochheiser K, Wiede F, Wagner T, Freestone D, Enders MH, Olshansky M, et al. Ptpn2 and KLRG1 regulate the generation and function of tissue-resident memory CD8+ T cells in skin. J Exp Med. 2021;218(6).

31. Shu Y, Ma X, Chen C, Wang Y, Sun X, Zhang L, et al. Myelin oligodendrocyte glycoprotein-associated disease is associated with BANK1, RNASE T2 and TNIP1 polymorphisms. J Neuroimmunol. 2022;372:577937.

32. Li J, Zaslavsky M, Su Y, Guo J, Sikora MJ, van Unen V, et al. KIR(+)CD8(+) T cells suppress pathogenic T cells and are active in autoimmune diseases and COVID-19. Science. 2022;376(6590):eabi9591.

33. Korner C, Altfeld M. Role of KIR3DS1 in human diseases. Front Immunol. 2012;3:326.

34. Sekine T, Perez-Potti A, Nguyen S, Gorin JB, Wu VH, Gostick E, et al. TOX is expressed by exhausted and polyfunctional human effector memory CD8(+) T cells. Sci Immunol. 2020;5(49).

35. Frieser D, Pignata A, Khajavi L, Shlesinger D, Gonzalez-Fierro C, Nguyen XH, et al. Tissue-resident CD8(+) T cells drive compartmentalized and chronic autoimmune damage against CNS neurons. Sci Transl Med. 2022;14(640):eabl6157.

36. Colamatteo A, Maggioli E, Azevedo Loiola R, Hamid Sheikh M, Cali G, Bruzzese D, et al. Reduced Annexin A1 Expression Associates with Disease Severity and Inflammation in Multiple Sclerosis Patients. J Immunol. 2019;203(7):1753–65.

37. Fernando MM, Stevens CR, Walsh EC, De Jager PL, Goyette P, Plenge RM, et al. Defining the role of the MHC in autoimmunity: a review and pooled analysis. PLoS Genet. 2008;4(4):e1000024.

38. Muniz-Castrillo S, Vogrig A, Honnorat J. Associations between HLA and autoimmune neurological diseases with autoantibodies. Auto Immun Highlights. 2020;11(1):2.

39. Wiendl H, Gross CC, Bauer J, Merkler D, Prat A, Liblau R. Fundamental mechanistic insights from rare but paradigmatic neuroimmunological diseases. Nat Rev Neurol. 2021;17(7):433–47.

34. Depuydt, M.A.C., Schaftenaar, F.H., Prange, K.H.M. et al. Single-cell T cell receptor sequencing of paired human atherosclerotic plaques and blood reveals autoimmune-like features of expanded effector T cells. Nat Cardiovasc Res 2, 112–125 (2023).

35. Li Y, Li B, You Z, Zhang J, Wei Y, Li Y, Chen Y, Huang B, Wang Q, Miao Q, Peng Y, Fang J, Gershwin ME, Tang R, Greenberg SA, Ma X. Cytotoxic KLRG1 expressing lymphocytes invade portal tracts in primary biliary cholangitis. J Autoimmun. 2019 Sep;103:102293

## References

1. Perriot S, Mathias A, Perriard G, Canales M, Jonkmans N, Merienne N, et al. Human Induced Pluripotent Stem Cell-Derived Astrocytes Are Differentially Activated by Multiple Sclerosis-Associated Cytokines. Stem Cell Reports. 2018;11(5):1199–210.

2. Perriot S, Canales M, Mathias A, Du Pasquier R. Generation of transgene-free human induced pluripotent stem cells from erythroblasts in feeder-free conditions. STAR Protoc. 2022;3(3):101620.

3. Perriot S, Canales M, Mathias A, Du Pasquier R. Differentiation of functional astrocytes from human-induced pluripotent stem cells in chemically defined media. STAR Protoc. 2021;2(4):100902.

4. Ho SM, Hartley BJ, Tcw J, Beaumont M, Stafford K, Slesinger PA, et al. Rapid Ngn2-induction of excitatory neurons from hiPSC-derived neural progenitor cells. Methods. 2016;101:113–24.

5. Callegari I, Oechtering J, Schneider M, Perriot S, Mathias A, Voortman MM, et al. Cell-binding IgM in CSF is distinctive of multiple sclerosis and targets the iron transporter SCARA5. Brain. 2024;147(3):839–48.

6. Mathias A, Perriot S, Jones S, Canales M, Bernard-Valnet R, Gimenez M, et al. Human stem cell derived neurons and astrocytes to detect auto-reactive IgG in neurological diseases. bioRxiv. 2024:2024.02.26.582006.

7. Genolet R, Bobisse S, Chiffelle J, Arnaud M, Petremand R, Queiroz L, et al. TCR sequencing and cloning methods for repertoire analysis and isolation of tumor-reactive TCRs. Cell Rep Methods. 2023;3(4):100459.

8. Chiffelle J, Barras D, Pétremand R, Orcurto A, Bobisse S, Arnaud M, et al. Tumor-reactive clonotype dynamics underlying clinical response to TIL therapy in melanoma. bioRxiv. 2023:2023.07.21.544585.

9. Borcherding N, Bormann NL, Kraus G. scRepertoire: An R-based toolkit for single-cell immune receptor analysis. F1000Res. 2020;9:47.

10. Hao Y, Stuart T, Kowalski MH, Choudhary S, Hoffman P, Hartman A, et al. Dictionary learning for integrative, multimodal and scalable single-cell analysis. Nat Biotechnol. 2024;42(2):293–304.

